# TraC of the conjugative plasmid pKM101 is secreted as a monomer in outer membrane vesicles and stimulates conjugation

**DOI:** 10.1101/2022.07.29.501992

**Authors:** Jaafar Amro, Christian Baron

## Abstract

Gram-negative bacteria use membrane-bound type IV secretion systems to assemble pili on the cell surface, followed by cell-cell contact with recipient cells and transfer of plasmid DNA. The process at the cell-cell contact stage of conjugative DNA transfer is not well understood. We here present a biochemical and genetic characterization of the TraC protein that is a minor component of the pili determined by the IncN plasmid pKM101 from *Escherichia coli*. The cellular and secreted forms of TraC are monomers, TraC preferentially localizes at the cell poles and it is also detected in extracellular membrane vesicles. Purified TraC does not impact the infection with bacteriophages, but we detect binding of TraC to recipient cells and partial complementation of a *traC* deletion strain by the addition of purified TraC. These results suggest that the protein contributes to conjugation at the cell-cell contact stage.

## Introduction

Type IV secretion systems (T4SSs) are transmembrane nanomachines present in Gram-negative and Gram-positive bacteria (1). T4SSs mediate bacterial plasmid transfer by conjugation, and they are responsible for effector protein secretion and pathogenicity in many pathogens of humans, animals and plants (2, 3). T4SSs are a diverse group of protein complexes, and their most conserved form in Gram-negative bacteria is composed of 12 proteins that form a transmembrane channel and an extracellular appendage, called the pilus (4). Based on sequence similarities the 12 subunit proteins are homologs of VirB1-11 and VirD4 based on the most studied T4SS of the plant pathogen *Agrobacterium tumefaciens* (5). During conjugation genetic material will be transferred from a donor to a recipient cell and this process requires cell-to-cell contact and possibly a membrane fusion event (6, 7). The transfer of genetic material enables bacteria to adapt to changing environmental conditions including exposure to antibiotics. The transfer for antibiotic resistance genes represents a global threat for public health (8-10).

The general architecture of T4SS is well documented. They comprise an outer membrane core complex and an inner membrane complex that are connected by a stalk forming the translocation channel (4, 11-13). The cell surface-exposed pilus comprises the major pilin VirB2 and the minor pilus tip protein VirB5 that is believed to be involved in the recognition and adhesion to recipient bacteria during conjugation (14-18). T4SS also represent the receptor for certain bacteriophages specific to bacteria expressing conjugative plasmids (19-22). VirB5 proteins are exposed on the surface and at the tips of pili and they may be adhesins that contact recipient cells. However, there is little direct evidence to support this notion except in case of the VirB5-homologous CagL protein from the human pathogen *H. pylori*, that binds to and activates integrin α_5_β_1_ receptors on gastric epithelial cells (15, 16, 23-25). The VirB5-like protein TraC from the conjugative plasmid pKM101 from *Escherichia coli* is translocated to the periplasm, followed by cleavage of its signal peptide before being routed to the cell surface using a mechanism probably involving the Bam proteins (23, 26, 27). TraC is a minor component of the T4SS-determined pilus and it is also secreted into the extracellular medium as a soluble protein in *E. coli* (26).

To understand the mechanistic contribution of TraC to conjugation we purified and characterized the cellular and secreted forms and found that the protein is a monomer in both locations. Analysis of the subcellular localization of TraC by immunofluorescence showed that it localizes at the poles and at the cell perimeter. TraC is also detected in outer membrane vesicles purified from the culture supernatant. Extracellular TraC does not impact phage infection, but it binds to recipient cells and partly restores conjugation of a Δ*traC* mutant pKM101 suggesting that it plays a role at the cell-cell contact stage of the conjugation process.

## Results

### Expression and purification of TraC from different subcellular localizations

To study TraC and its interactions with other proteins in different subcellular localisations, we overexpressed and purified a C-terminally Strep-tagged version (TraC_Strep_) from *E. coli* in the presence or absence of pKM101. TraC_Strep_ was expressed from plasmid pTrc200 and complementation of pKM101Δ*traC*1134 carrying a non-polar transposon insertion in the *traC* gene (28) showed that the modified protein was fully functional as measured by plasmid conjugation and phage infection assays (**suppl. Fig. 1**). We isolated TraC_Strep_ from the cell supernatant (SN), from the membranes (M) and from the soluble fraction of cell lysates (S), followed by SDS-PAGE, silver staining and Western blotting (**Fig. 1**). In addition to detecting TraC_Strep_ this analysis revealed the presence of other proteins that were identified by mass spectrometry. OmpA (∼50kDa) was detected in the membrane fraction, the lipoprotein LPP (∼10 kDa) in the fraction of secreted TraC_Strep_ and the biotin carboxyl carrier protein (BCCP) was detected in the soluble fractions (∼13 kDa) (**suppl. Fig. 2**). We obtained similar results when TraC was purified from *E. coli* with or without pKM101.

**Figure 1.**
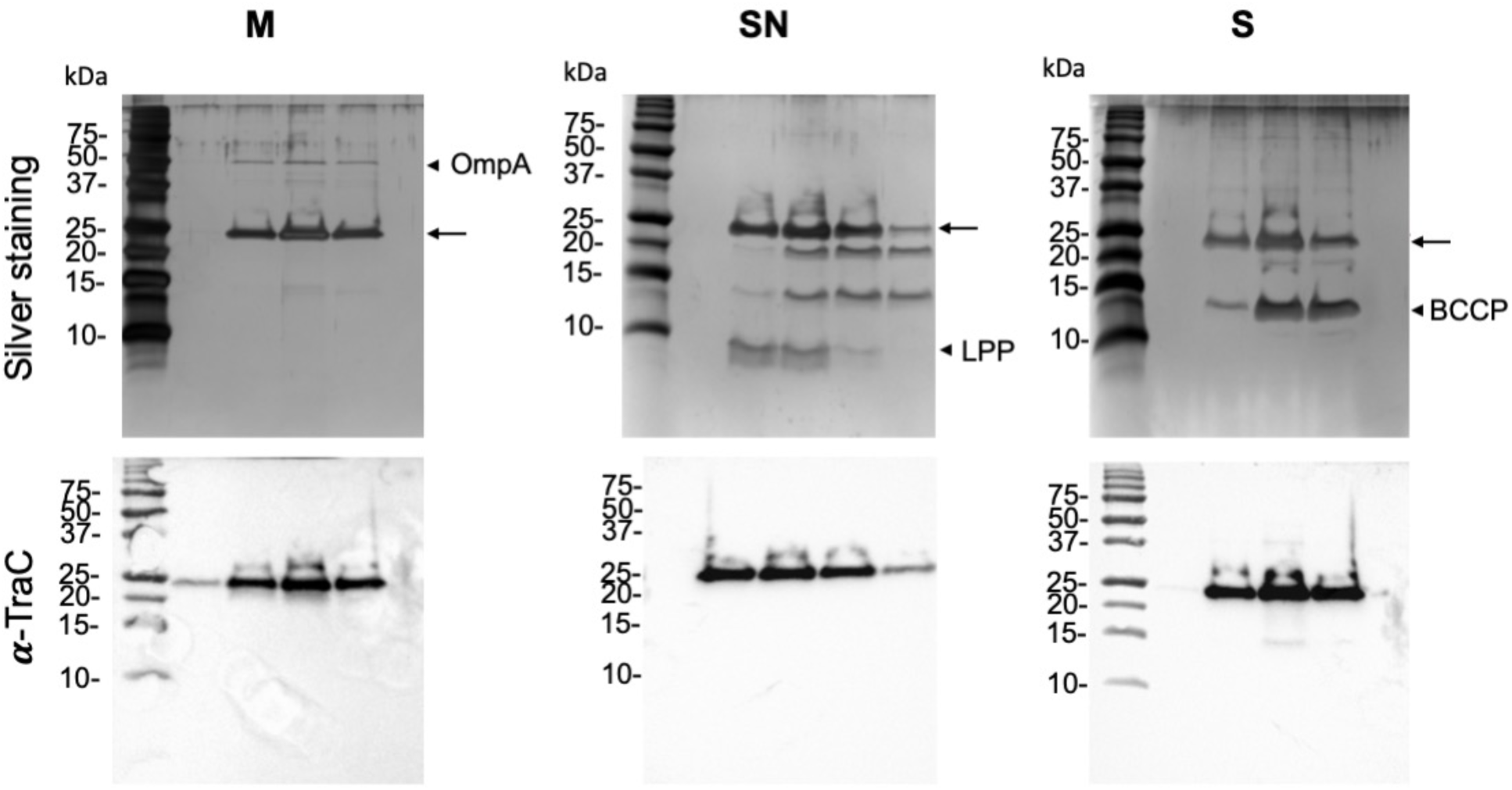
Tricine-SDS-PAGE analysis of the elution fractions after strep-column affinity purification. (Upper panel) silver staining, (lower panel) western blot analysis with TraC-specific antiserum. Arrows indicate the molecular weight of TraC. S: soluble, SN: secreted, M: membrane, MW: molecular weight marker.

**Figure 2.**
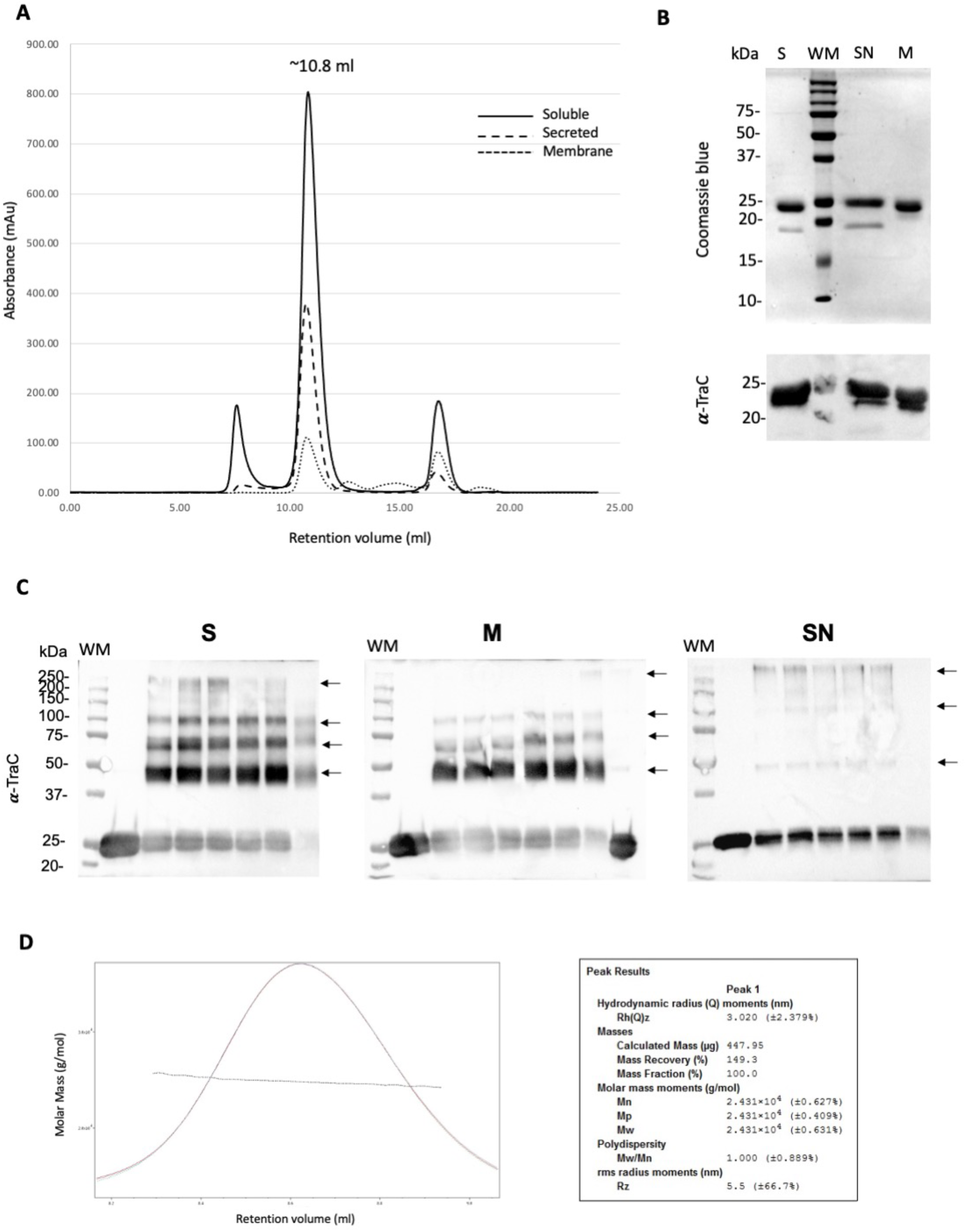
Purification of soluble, membrane-bound and secreted TraC_Strep_. (A) Elution profile of TraC_Strep_ with size exclusion chromatography. (B) (upper panel) SDS-PAGE analysis of the purified TraC_Strep_, (lower panel) western blot analysis with TraC-specific antiserum. (C) Western blot analysis with TraC specific antiserum of the purified periplasmic, membrane and secreted TraC_Strep_ in the absence and presence of increasing concentrations (0.4 – 4 mM) of the cross-linking agent DSS. Arrows indicate higher molecular-weight complexes formed after cross-linking. (D) Analysis of the oligomerization state of TraC. Elution profile of the TraC (left) is shown with the molecular weight estimated by MALS (right). S: soluble, SN: secreted, M: membrane, MW: molecular weight marker.

### TraC is primarily a monomer

To further characterize TraC_Strep_ from the different compartments, the fractions from affinity purification were pooled and concentrated, followed by size exclusion chromatography over a S75 column as additional purification step. TraC_Strep_ eluted at the same position from the column (∼10.8 ml) in all cases and it was most concentrated when it was purified from the soluble fraction of cell lysates (**Fig. 2A and B**). Interestingly, the elution profile was identical when the purification was conducted from cells with or without pKM101 indicating that TraC_Strep_ does not interact strongly with other T4SS components (data not shown). The retention volume corresponds to a molecular weight of ∼37 kDa, which is larger than the predicted size of the TraC monomer (∼24 kDa).

To characterize the oligomeric state of purified TraC_Strep_ from the SN, M and S fractions, we added increasing concentrations of the homo-bifunctional cross-linking agent disuccinimidyl suberate (DSS), followed by SDS-PAGE and western blotting. This analysis showed the formation of higher molecular mass complexes that may correspond to dimers as well as multimers of TraC or to complexes with other proteins (**Fig. 2C**). The crosslinking patterns of the M and S fractions were similar, but we observed a lower amount of higher molecular mass complexes in the SN fractions of secreted TraC_Strep_ indicating a lower degree of complex formation.

To precisely calculate the molecular weight of purified TraC_Strep_ we used SEC-MALS (size exclusion chromatography-multi angle light scattering) and this analysis revealed that the protein is monodisperse with a molecular mass of 24.3 kDa (**Fig. 2D**). We conclude that TraC is primarily a monomer that may undergo transient interactions with itself (dimerization) or with other proteins in the different subcellular compartments.

### TraC accumulates at the cell poles

Next, we used super resolution microscopy (structured illumination microscopy, SIM) and immunofluorescence microscopy to localize TraC in *E. coli*. To ensure the biological significance of our findings we expressed C-terminally Flag-tagged TraC (TraC_Flag_) in a pKM101Δ*traC*1134 mutant strain using the tightly controlled pBAD promoter. The level of expression was modulated by varying the concentration of the inducer arabinose to correspond to that of the endogenous TraC expressed from pKM101 and 10^−3^ % (66.5 μM arabinose) resulted in comparable levels of expression (**suppl.Fig. 3A**). We next tested plasmid conjugation (**suppl. Fig. 3B**) and phage infection (**suppl. Fig. 3C**) showing that TraC_Flag_ is fully functional under these conditions. *E. coli* cells were then permeabilized for the detection of cell-bound TraC with specific antisera and SIM localized the protein at the cell perimeter and at the cell poles (**Fig. 3A**). We observed the same polar localization in cells expressing, or not, a functional T4SS. We also incubated the donor cells (cells expressing TraC_Flag_ + T4SS) with plasmid-free recipient cells to assess whether there are changes of the localization of TraC during conjugation. We did not detect any difference in the TraC localization pattern in the presence of recipient cells (data not shown).

**Figure 3.**
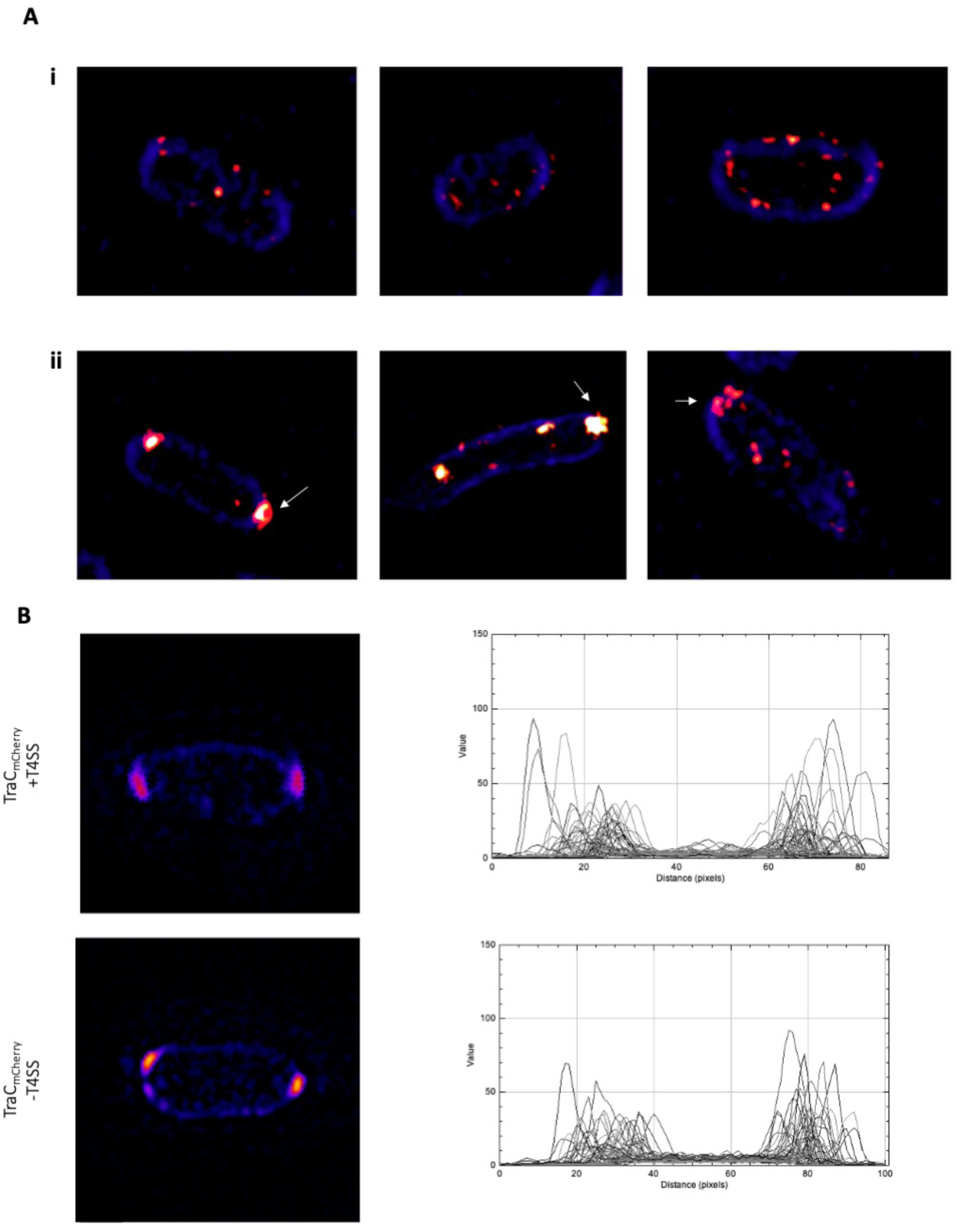
Cellular localization of TraC. (A) Cell producing TraC_Flag_ in the presence of the pKM101 T4SS. TraC_Flag_ foci were detected by reactivity with anti-Flag mouse primary and AlexaFluor 594-conjugated goat anti-mouse secondary antibody (red) at the perimeter (i) and at the pole (ii) of the cell. Cell membrane was visualized by staining with TMA-DPH (blue). White arrows indicate polar accumulation of TraC_Flag_. (B) Cell producing TraC_mCherry_ in the presence, or absence, of the pKM101 T4SS. TraC_mCherry_ (in red) were detected by structural illumination microscopy using mCherry specific filters (left panel). Quantification of the fluorescence intensity across 50 cells showing preferentially polar localization of TraC (right panel).

As a complementary approach, we engineered a C-terminal fusion of TraC to the fluorescent protein mCherry that was equally expressed at levels corresponding to TraC in pKM101 when induced with 10^−4^% arabinose (**suppl. Fig. 4A**). TraC-mCherry was partially functional during conjugation (∼0.2% of wild type TraC, **suppl. Fig. 4B)**. Similar to TraC_Flag_ we observed mostly polar TraC-mCherry (**Fig. 3B)** confirming the primary subcellular localization of the protein.

**Figure 4.**
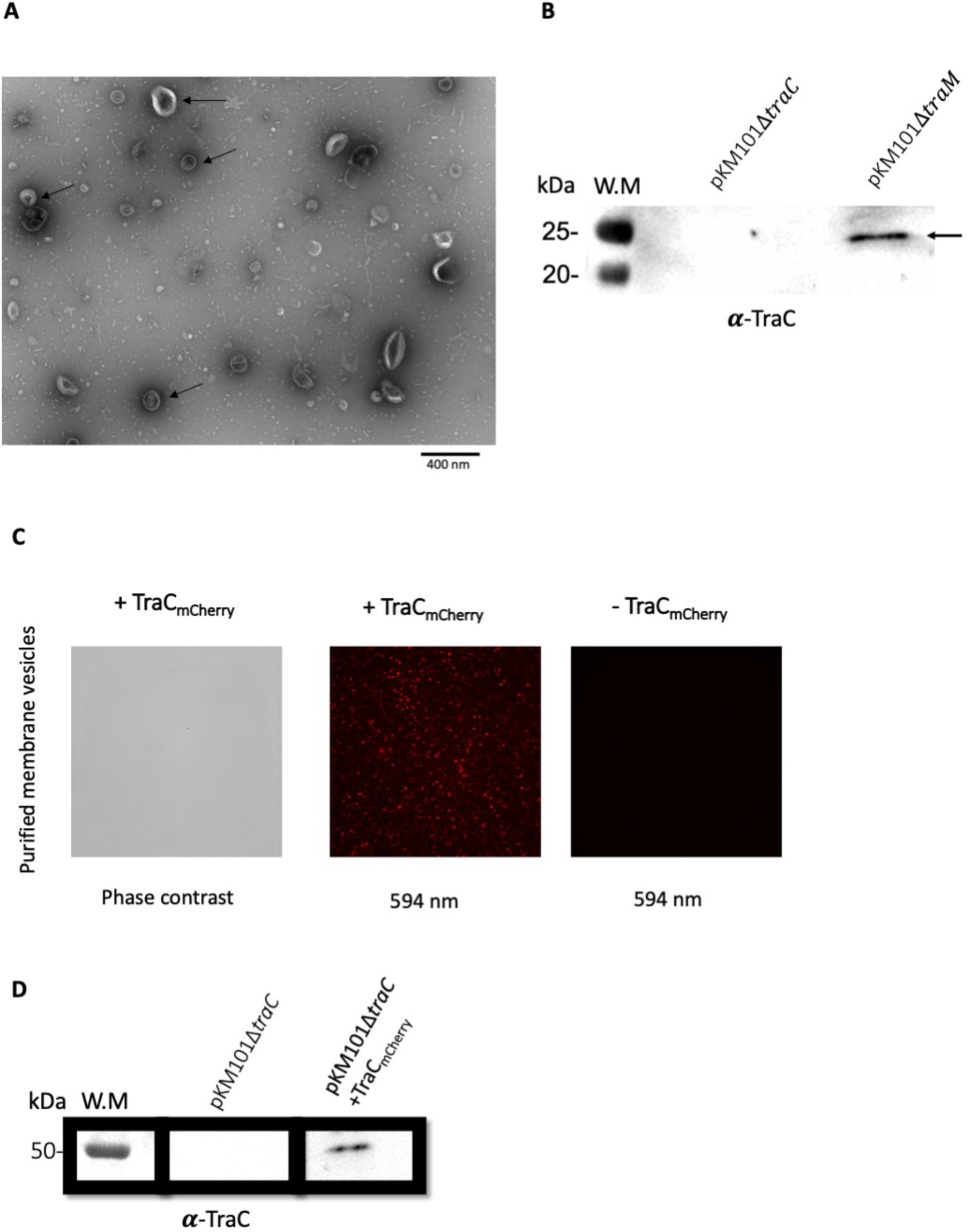
Detection of TraC in purified outer membrane vesicles (OMV). (A) Purified OMV under negative staining electron microscopy. Arrows point to OMV in different sizes. (B) western blot analysis using TraC-specific antiserum, of the purified OMV from a TraC-deficient and TraM-deficient cells culture supernatant. Arrow indicate the molecular weight of TraC. (C) Purified OMV under fluorescence microscopy from TraC_mCherry_ producing cells. (D) western blot analysis using TraC-specific antiserum, of the purified OMV from culture supernatant of TraC-deficient and TraC_mCherry_ producing cells.

### TraC is secreted with membrane vesicles

Outer membrane vesicles (OMV) exist in almost all Gram-negative bacteria, and play an important role in many bacterial functions, including cell communication, biofilm formation and pathogenicity (29). Since extracellular secretion of TraC was previously reported (23, 26) we next assessed whether it is secreted as part of OMVs. To this effect, we purified OMVs from the supernatant of *E. coli* carrying pKM101Δ*traM*363 that do not produce pili since they lack the major pilus component TraM. TraC was detected by SDS-PAGE and western blotting in the OMV fraction showing spherical particles of different sizes visualized by electron microscopy (**Fig. 4A and B**). As an independent approach to confirm the presence of TraC in OMV we purified OMVs from *E. coli* strain pKM101Δ*traC*1134 expressing TraC fused to mCherry at its C-terminus. Using fluorescence microscopy we detected mCherry in the OMV sample and western blotting confirmed the presence of the fusion protein (**Fig. 4C and D**) confirming that TraC is secreted as part of OMVs.

### Extracellular TraC does not affect IncN specific phage infection

Ike and PRDI are two phages that recognize and infect bacteria that carry IncN group conjugative plasmids (30). Ike attaches to the pilus tip before injecting its ssDNA into the bacteria, and PRDI attaches to the cell surface in a T4SS-dependent manner (27, 30). Previous studies showed that a point mutation in TraC (Val144) renders the cell resistant to PRDI without affecting Ike infection (27). These results suggest that TraC may be part of the host recognition site for phage infection. Therefore, addition of soluble TraC may decrease infection by blocking the phage receptors. We tested this hypothesis by mixing Ike and PRDI phages with 5 μg of purified TraC before the infection. We did not detect any effect of soluble TraC on the infection rate (**Fig. 5A**). We also tested whether extracellular TraC renders pKM101Δ*traC*1134-carrying cells susceptible to Ike and PRDI infection (27). To this effect, we added 5 μg of soluble TraC to pKM101Δ*traC*1134-carrying cells, but we did not detect any plaque formation (**Fig. 5A**). We conclude that extracellular TraC does not block infection and does not restore the sensitivity of TraC-deficient cells toward Ike or PRD1. Finally, we used crosslinking to detect whether there are interactions between TraC and the phages. Indeed, in the presence of the crosslinking agent disuccinimidyl suberate (DSS), we identified cross-linking products indicating a potential interaction between TraC and PRD1 proteins, but not with Ike (**Fig. 5B**). The apparition of distinct crosslinking products, e.g. of ∼150 kDa, may reflect this interaction. The receptor binding protein P2 of PRD1 has a size of 64 kDa and it forms dimers (31, 32). It will be interesting to further study the composition of the observed crosslinking products to understand the phage-cell interaction.

**Figure 5.**
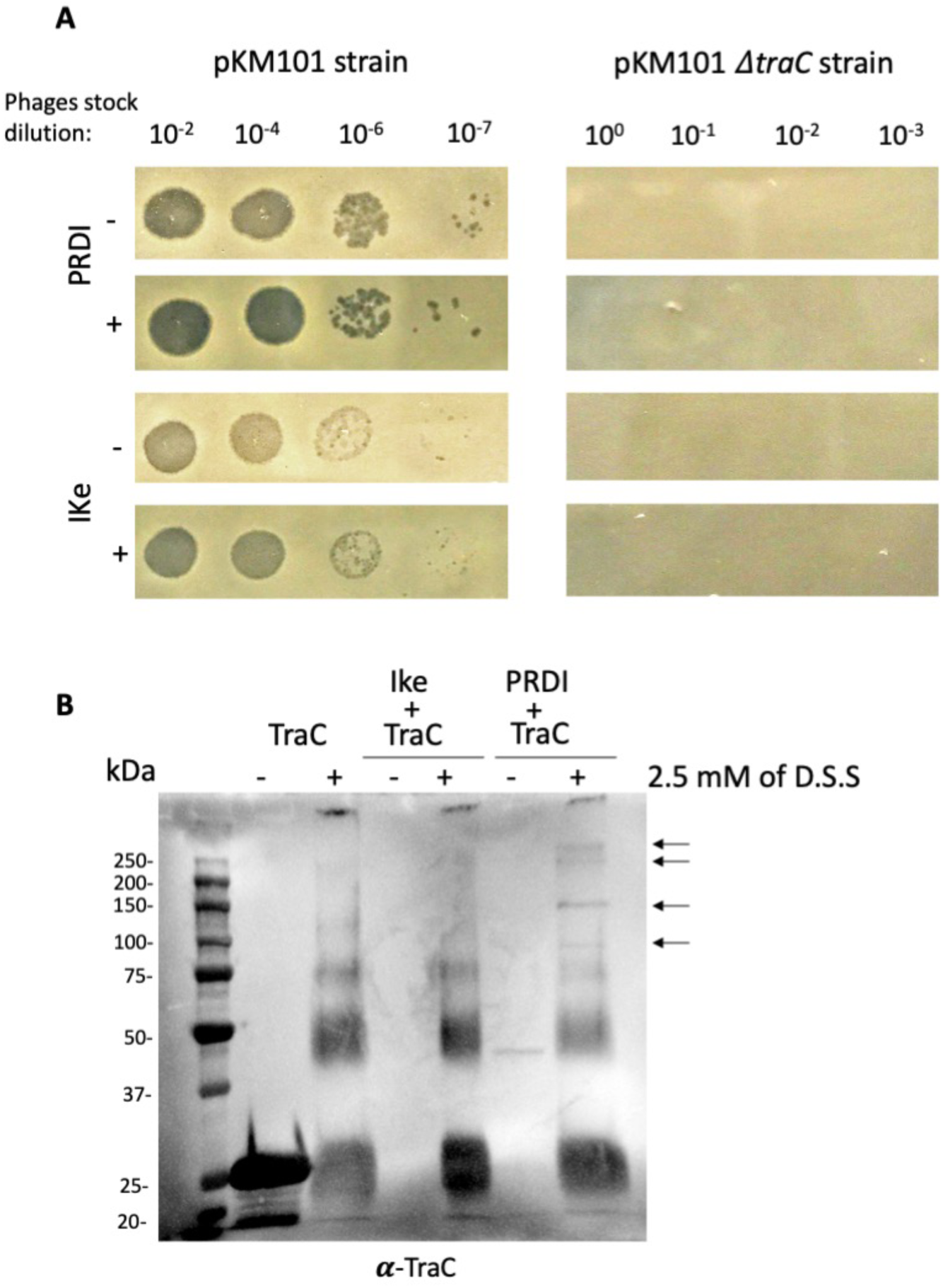
Effect of extracellular TraC on IncN specific phages infection. (A) Phage infection rates monitored using a TraC-deficient cells producing increasing amount of TraC expressed under *ara* pBAD promoter. (B) The effect of 5 μg of purified TraC on pKM101 or TraC-deficient cells sensitivity toward a serial dilution of Ike and PRDI phages. (+) and (-) indicate the presence or absence of purified TraC during the infection. (C) Western blot analysis using TraC-specific antiserum, of Ike and PRDI interaction with 5 μg of purified TraC, in the absence or presence of 2.5 mM of DSS. Arrows indicate a high molecular weight complex formed after crosslinking.

### Purified TraC partially restores the transfer of pKM101ΔtraC plasmid

Previous studies showed that a donor bacterium carrying pKM101Δ*traC*1134 can be partially complemented by a helper cell expressing a functional T4SS (26, 28). Here, we tested the effect of purified TraC on plasmid transfer from pKM101Δ*traC*1134 donor cells. 5 μg of purified TraC were mixed with donor and recipient cells on a solid surface (∼10^8^ copies of TraC per cell), followed by quantification of conjugative transfer. The results show a significant increase of the transconjugant numbers in the presence of extracellular TraC (∼200 fold) (**Fig. 6A**). The addition of purified TraC does not have a measurable impact on the conjugation rate of *E. coli* carrying the pKM101 donor or the R388 plasmid control. The partial restoration of plasmid transfer suggests an extracellular role of secreted TraC during conjugation and it may interact with recipient cells. To test this possibility, we mixed purified TraC with recipient cells and indeed TraC co-sedimented with recipient cells indicating that it may bind to recipients (**Fig. 6B**). Future studies will show whether this observation reflects a specific interaction and whether there is a specific receptor on recipient cells.

**Figure 6.**
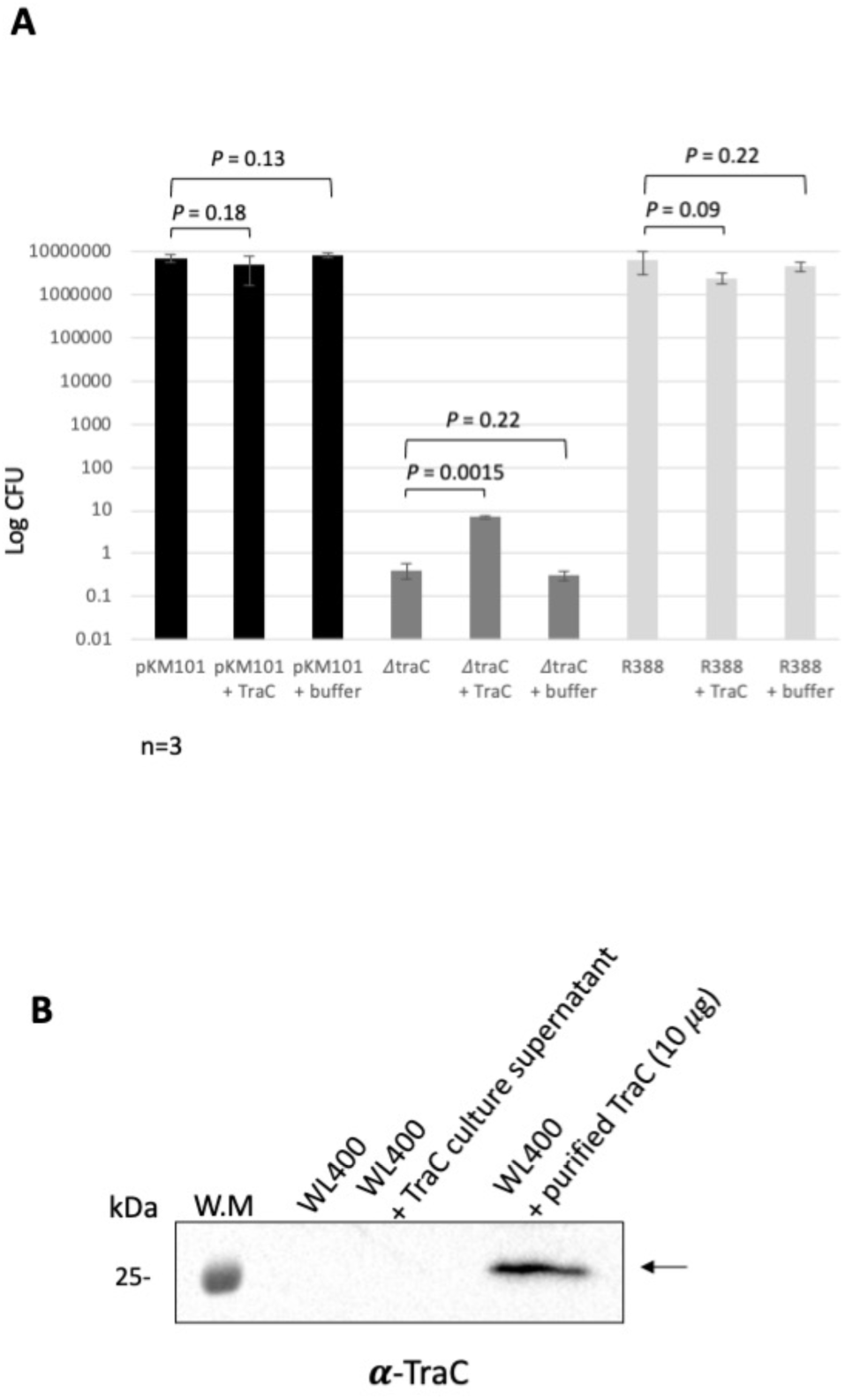
Effect of extracellular TraC on conjugation efficacy. (A) The effect of 5 μg of purified TraC, or buffer alone, on the plasmid transfer rate during conjugation for wild type pKM101, TraC-deficient strain, and R388 plasmid, containing strain. Values and S.D. (*error bars*) were calculated from three independent experiments. (B) Western blot analysis using TraC-specific antiserum, for the interaction between purified TraC (10 μg) or secreted TraC from 1 ml culture, with the recipient cell WL400. Arrow indicates the molecular weight of TraC.

## Discussion

TraC is known to locate in various cellular compartments, e.g. in the membrane, in the periplasm and attached to the pilus. TraC is also exposed to the surface and secreted to the extracellular medium in a soluble form (23, 26). We purified cell-bound and secreted Strep-tagged TraC and we detected other proteins that co-eluted after affinity purification. BCCP is the only biotinylated protein in *E. coli*, and it is known to co-purify as a contaminant when using the Strep-tag affinity columns (33). This probably explains why it co-eluted with TraC from the soluble (S) fraction. The other co-eluted proteins may represent specific interaction with tagged TraC. In a previous study, crosslinking showed an interaction between TraC and many outer membrane proteins including Bam proteins and OmpA as we have found here in the case of membrane-bound TraC (M fraction) (23). The major outer membrane protein LPP is the most abundant cellular protein and it is present in the periplasm and exposed to the cell surface (34). LPP co-eluted only with the secreted form of TraC suggesting that it may undergo a specific interaction during the secretion process. SEC-MALS showed that TraC is a monomer when purified from different subcellular localizations (S, M and SN), despite the fact that crosslinking showed formation of higher molecular weight complexes in the presence of DSS. The crosslinking results could be interpreted as evidence for transient non-specific protein-protein interactions between TraC and other proteins such as OmpA and LPP.

Several T4SS proteins have been localized by immuno-localization at the cell pole (35-38). In *A. tumefaciens* the polar localization of VirB5 requires the presence of VirB8 and VirB9 (36), and VirB5-like proteins are also suggested to be present as in the inner membrane complex of the T4SS (4, 13). Here, we used two complementary microscopic approaches to localize TraC in permeabilized cells demonstrating accumulation at the cell perimeter and at the pole. The localization was independent of other T4SS components. These results indicate that the polar accumulation of TraC reflects localization of TraC in the periplasm or in the inner membrane. In contrast, a previous study of non-permeabilized cells by immunofluorescence detected TraC as distinct foci on the cell surface (23). These results suggest two different routes for the trafficking of TraC. After its translocation to the periplasm by the signal peptide, TraC is directed to the cell surface and secreted into the extracellular medium in a T4SS-independent fashion. Alternatively, TraC accumulates at the cell pole where it may integrate the T4SS. It is noteworthy that the T-pilus in *A. tumefaciens* assembles at the cell pole probably reflecting a T4SS assembly site (39).

The conjugative phenotype of *E. coli* expressing a TraC-deficient T4SS can be partially restored by the presence of a helper cell producing a functional pKM101 T4SS (26, 28). We here tested the effect of purified TraC during conjugation and phage infection of a TraC-deficient strain (carrying pKM101Δ*traC*1134). We detected a significant 200-fold increase of plasmid transfer during conjugation, but no effect on Ike and PRD1 phage infection. Interestingly, we showed that TraC may interact with the recipient cells, and further experiments are needed to identify whether this interaction is specific. Thus, extracellular TraC could promote cell-to-cell contact during conjugation by promoting an interaction with the recipient cell. This is in agreement with previous studies showing a role of VirB5-like proteins as adhesins during conjugation and host cell infection (15, 16, 23-25). The detection of TraC in purified OMV raised the possibility that neighboring bacteria can share T4SS subunits to complement deficiencies and this may be the basis for extracellular complementation. It will be interesting to assess whether other T4SS subunits are present in purified OMV, and whether these vesicles influence the efficacy of conjugation and of phage infection.

## Acknowledgements

This work was supported by grants from the Natural Sciences and Engineering Research Council (NSERC, #RGPIN-2017-05123) and the Canadian Institutes of Health Research (CIHR, #398288 and #274108) to C.B. We grateful to Dr. Aurélien Fouillien at the Université de Montréal EM facility in the Faculty of Dentistry for technical support and assistance. We also thank Dr. Nicolas Stifani and Dr. Normand Cyr (Department of Biochemistry and Molecular Medicine) at the Université de Montréal Faculty of Medicine for technical support and help with data analysis.

## Experimental procedures

### Strains and growth conditions

The strains used are listed in supplementary Table 1. *E. coli* FM433 and derivatives were grown in Luria-Bertani (LB) media supplemented with streptomycin (100 μg/ml), spectinomycin (100 μg/ml), ampicillin (100 μg/ml), chloramphenicol (34 μg/ml), for plasmid propagation or selection of transconjugants. For induction of the LacI-repressed *trc* promoter in pTrc200 constructs, isopropyl-β-D-thiogalactopyranoside (IPTG) was added to a final concentration of 0.5 mM.

L-arabinose was used for the induction of the *ara* pBAD promoter.

### Protein expression and purification

TraC was overproduce and purified as described elsewhere with minor modification (27, 40). *E. coli* strain BL21star (λDE3) harboring pTrcTraC_Strep_, was grown in LB supplemented with 50 μg/mL kanamycin and 50 μg/mL streptomycin. Overnight precultures in LB were used to inoculate 3 litres culture (37 °C) until they reached an OD_600_ of 0.6–0.8. Expression was induced by addition of 0.2 mM IPTG at 37 °C, and cultures were left incubated for 3 h. For purification, bacterial cells were harvested, and the obtained supernatant was purified using 0.22 μm filter and passed over a Strep-Tactin II column (GE Healthcare) to collect secreted TraC, and further eluted with 2.5 mM desthiobiotin. The harvested bacteria were resuspended in binding buffer (100 mM Tris-HCl pH8, 150 mM NaCl, 1 mM EDTA) with cOmplete Mini Protease Inhibitor mixture and DNase I at 100 μg/mL, and lysed twice using a One Shot cell disrupter (Constant Systems, Inc.) at 27 kpsi and 4 °C. Debris was removed by centrifugation twice at 15000 × *g* for 30 min at 4 °C, and the supernatant was retained. Pursuing ultracentrifugation at 250,000 × *g* for 1 h at 4 °C, to separate soluble and membrane bound TraC. The soluble fraction was passed over Strep-Tactin II column and soluble TraC was eluted with 2.5 mM desthiobiotin. Total membranes were collected and solubilized overnight at 4 °C with gentle stirring binding buffer with 0.1% (wt/vol) detergent DDM with cOmplete Mini Protease Inhibitor mixture. This material was then centrifuged for 1h at 35000g × *g* to collect DDM-solubilized TraC for purification over a Strep-Tactin II column and eluted using 2.5 mM desthiobiotin. For biochemical analysis, TraC was further purified in phosphate buffer (50 mM phosphate, 100 mM NaCl pH 7.4), by size exclusion chromatography (SEC) using a Superdex 75 column (GE Healthcare).

### Protein complex analysis

The oligomerization state of TraC was analyzed by SEC-MALS as described in ref. (40). SEC-MALS was realized with the use of an ÄKTAmicro system (GE Healthcare) coupled to a Dawn HELEOS II MALS detector and an OptiLab T-rEX online refractive index detector (Wyatt Technology). The absolute molecular mass was calculated by analyzing the scattering data using the ASTRA analysis software package, version 6.1.6.5 (Wyatt Technology). Protein samples were separated on a Superdex 75 10/300 increase SEC column (GE Healthcare) with a flow rate of 0.5 mL/min. BSA was used for calibration. A 0.1-mL sample of TraC at concentrations ranging between 1 and 10 mg/mL was injected and eluted in 50 mM sodium phosphate buffer (pH 7.4), 100 mM NaCl. The molecular mass of TraC was determined by the dual detection method implemented in the conjugated analysis mode of the ASTRA analysis software.

### Conjugation assay

Quantitation of conjugative DNA transfer were monitored as described elsewhere with minor modifications (26). Equal amounts of donor and recipient cells from a mid-log phase culture were harvested and resuspended with LB medium without antibiotics. 1 μl of recipient and donor cells were mixed on a prewarmed LB agar plate and incubated for 2 h at 37°C. L-arabinose was mixed with the LB agar to induce the expression of TraC variants from a pBAD plasmid containing strains, during conjugation. The spot was washed from the plate three times with 150 μl of LB medium. To quantitate conjugative transfer, dilutions were plated on LB media containing appropriate antibiotics for selection of plasmid-containing recipients.

### Phage infection assay

Ike and PRDI bacteriophages infection were performed as described in reference (20). Briefly, 200 μl of cells from mid-log phase culture, in the presence of inducers for TraC variants expression, were mixed with 3 ml of top agar and poured on LB plate containing appropriate antibiotic and inducer, and allowed to dry. Drops of 5 μl of serially diluted phages were spotted on the surface of the top agar, and the plates were examined for plaques formation after overnight incubation at 37°C.

### Cross-linking assay

Chemical cross-linking with DSS (Pierce) was performed as described elsewhere with minor modifications (41). Briefly, DSS was added in different concentrations (0.4 to 4 mM) to 10 μl of purified TraC, and the samples were incubated for 30 min at RT, followed by the addition of 1 volume of Laemmli sample buffer and analysis by SDS-PAGE and Western blotting. For glutaraldehyde, different concentrations (0.001 % to 0.2 %) were used, and the incubation time was reduced to 5 min at R.T. For intracellular cross-linking, 1 ml of cell culture (O.D ∼0.8) were centrifuged at 15000 g for 10 min and bacteria pellet was washed three times and incubated with 5 mM DSS for 30 min at RT, followed by the addition of 1 volume of Laemmli sample buffer and analysis by SDS-PAGE and Western blotting.

### Outer membrane vesicle purification

Cells from late log phase liquid culture were removed by centrifugation two times (10 000g, 20 min). The supernatant was filtered through 0.22 μM filter (Millipore) and OMV were collected by high-speed centrifugation at 100,000g for 1 hour. The obtained crude OMV were analyzed by western blotting using TraC specific antiserum and visualized by fluorescent or electron microscopy.

### Negative-staining electron microscopy

5 μl of purified OMV were deposit on negatively glow-discharged 300 mesh copper grid for 3 min and blotted using Whatman filter paper. Samples were washed three times with filtered water, followed by staining with 2% uranyl acetate solution for 45 seconds. The stain was then removed by blotting the grid with Whatman filter paper and air drying at RT. The samples were imaged using a FEI Tecnai T12 120 kV TEM equipped with Gatan 2K AMT camera, in the Université de Montréal EM facility.

### Fluorescence microscopy

For fluorescence microscopy, cells producing TraC-mCherry were induced with 10^−4^% of L-arabinose for 2h and fixed with 4% of paraformaldehyde. Cells were washed extensively, adhered to poly-lysine coated coverslip, and inverted onto a drop of ProLong Gold (Life Technologies). Immunolabeling was realized as described elsewhere with minor modification (42). cells producing TraC_Flag_ were induced with 10^−3^% of L-arabinose for 2h and fixed with 4% of paraformaldehyde. Cells were washed extensively and permeabilized with 0.1% Triton-X100, 100 μg/ml of lysozyme and 5mM of EDTA. After washing 3 times cells were blocked with 2% BSA for 1h at 37 followed by incubation with 1/100 dilution of anti-Flag monoclonal antibody (Cell Signaling) for 1.5h at RT. Cells were then washed 3 times and incubated with 1/1000 of monoclonal secondary antibody conjugated to Alexa Fluor 594 (Invitrogen) for 45 min at RT in the dark. Cell membranes were stained with 10 mM of TMA-DPH for 10 min at RT in the dark. Cells were washed, adhered to poly-lysine coated coverslip, and inverted onto a drop of ProLong Gold (Life Technologies). 5 μl of samples were spotted on a coverslip and immobilized by 3 μl of ProLong Diamond (Life Technologies, Mississauga, ON, Canada). Samples were examined at room temperature with a ×63/1.4 oil objective under structured illumination microscopy using an Elyra PS1 microscope (Carl Zeiss, Oberkochen, Germany). mCherry proteins and Alexa fluor 594 conjugated antibodies were excited at 561 nm, and emission around 610 nm was observed using a 14-bit electron-multiplying charge-coupled device (EMCCD) camera. The membrane dye TMA-DPH was excited at 405 nm, and the emission was observed at around 460 nm. Z-stack volumes were acquired using the SIM and reconstructed using Zen Black edition software. The fluorescence of entire bacteria and of each bacterial pole was quantified using ImageJ software.

## Supplementary data

**Supplementary Table 1.**
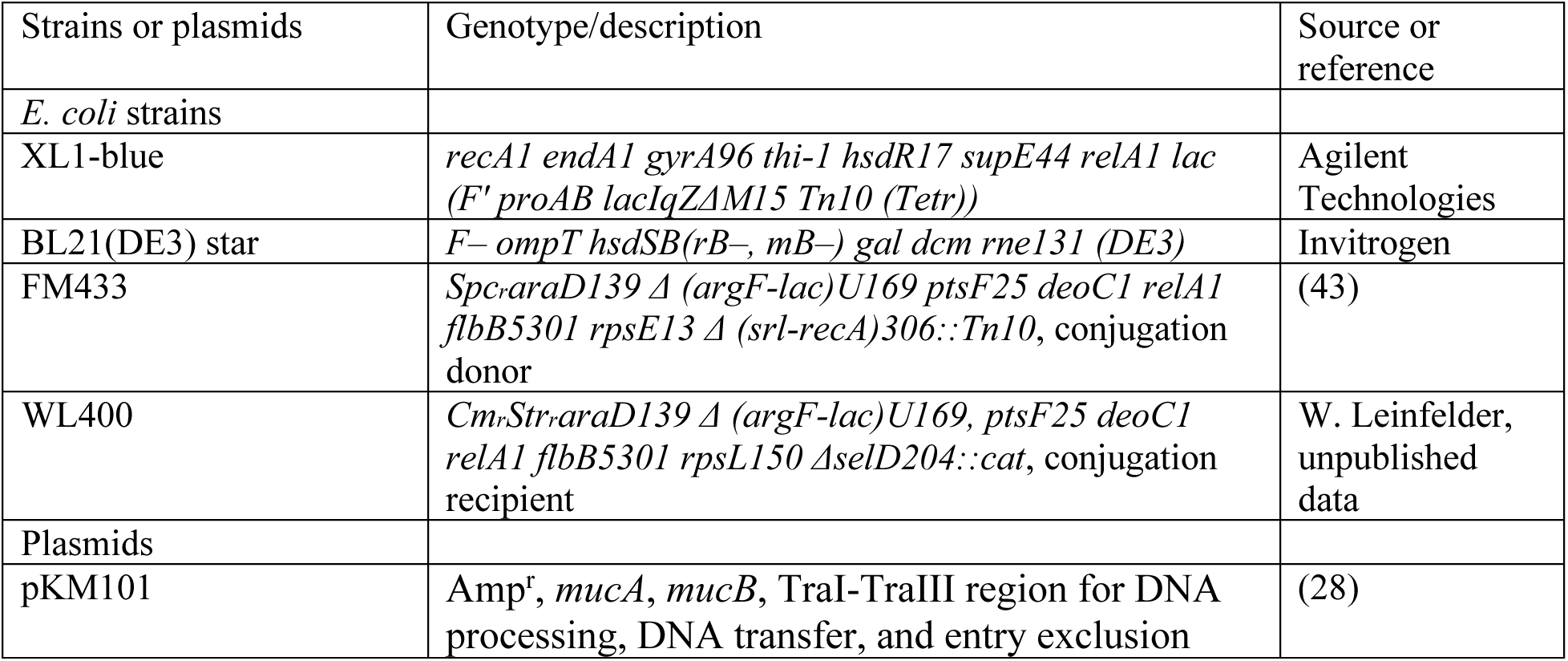
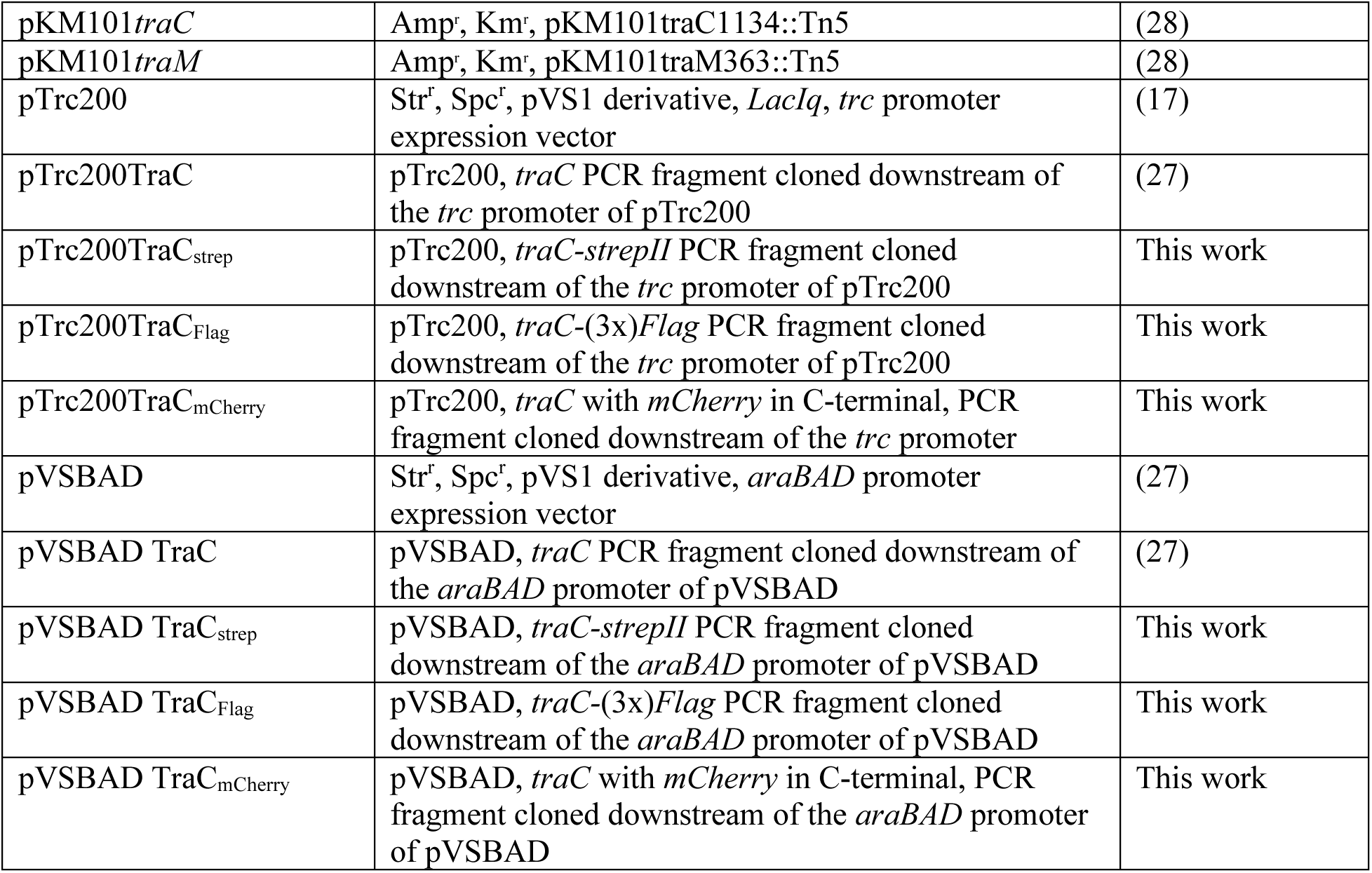
Bacterial strains and plasmids.

**Supplementary Figure 1.**
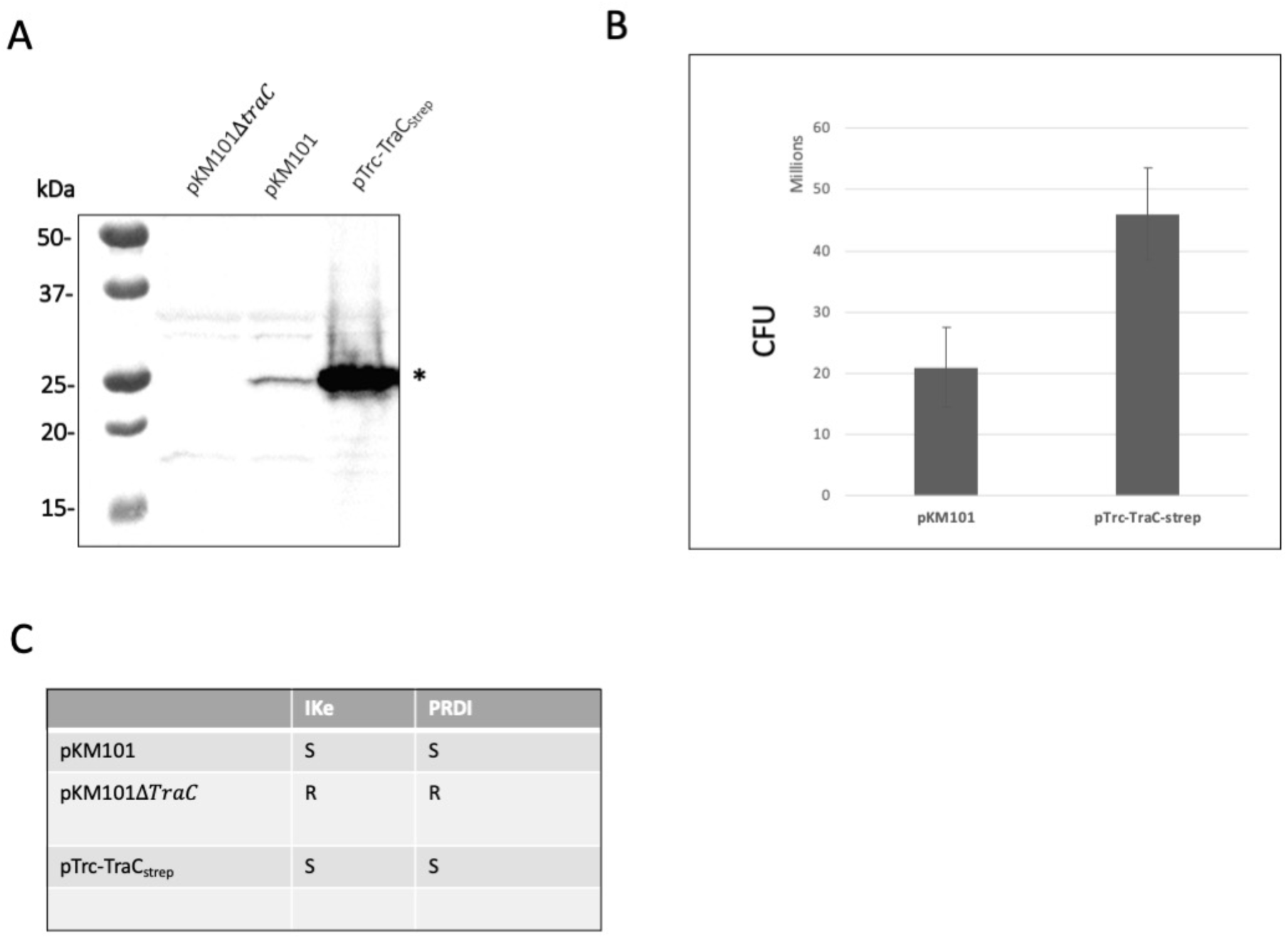
(A) Western blot analysis using TraC-specific antiserum, of TraC_strep_ expression level in total cell lysate under native promotor or pTrc200 promoter using 0.2 mM of IPTG. Asterisk (*) indicate TraC_Strep_. Functionality of TraC_strep_ was monitored by conjugation (B) and phage infection (C) using 0.2 mM IPTG. Values and S.D. (*error bars*) were calculated from three independent experiments.

**Supplementary Figure 2.**
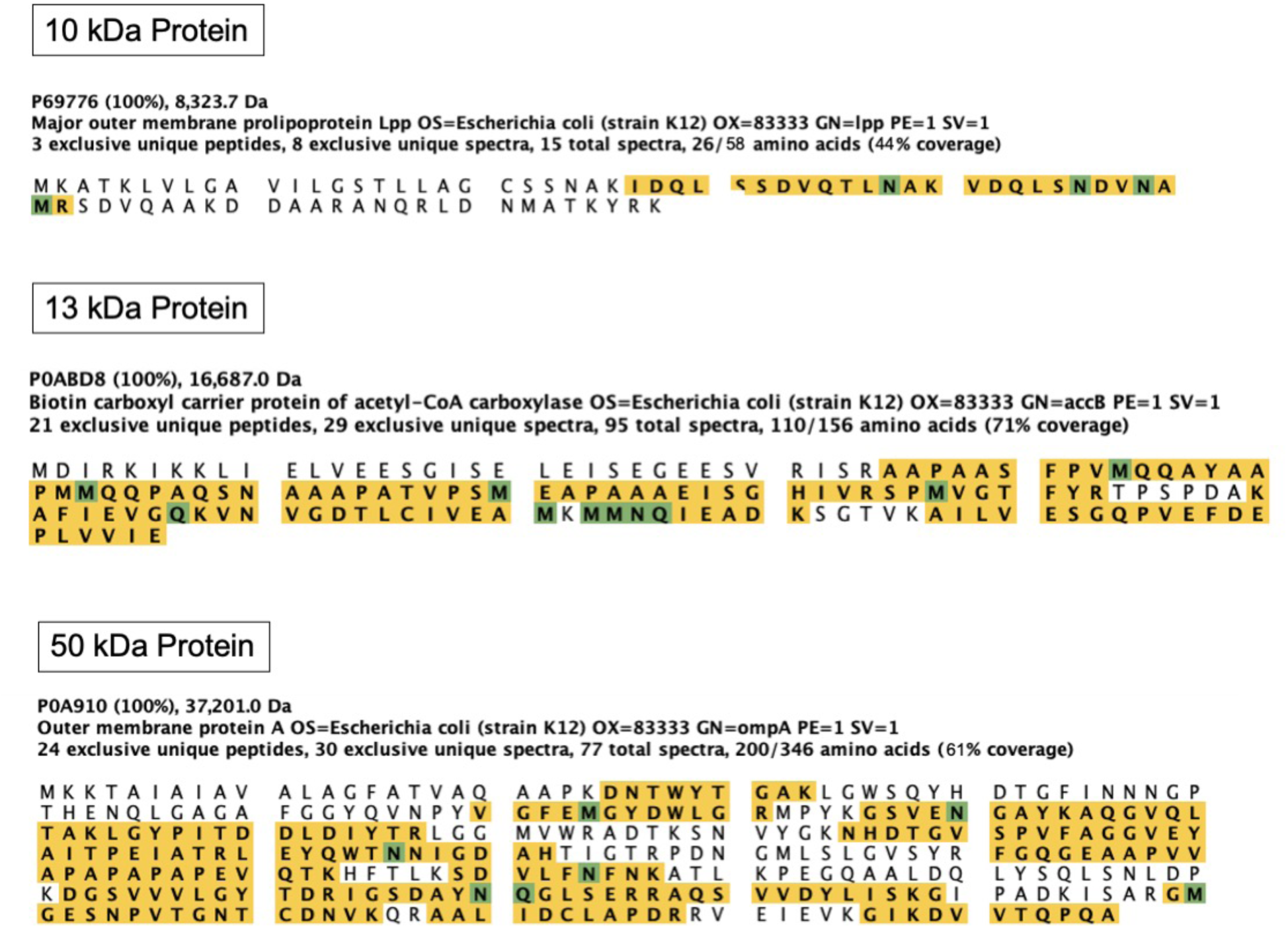
Mass spectrometry analysis of the co-eluted proteins shown in Figure 1. Proteins from gels were cut and analyzed by mass spectrometry. Identified peptides are highlighted in yellow.

**Supplementary Figure 3.**
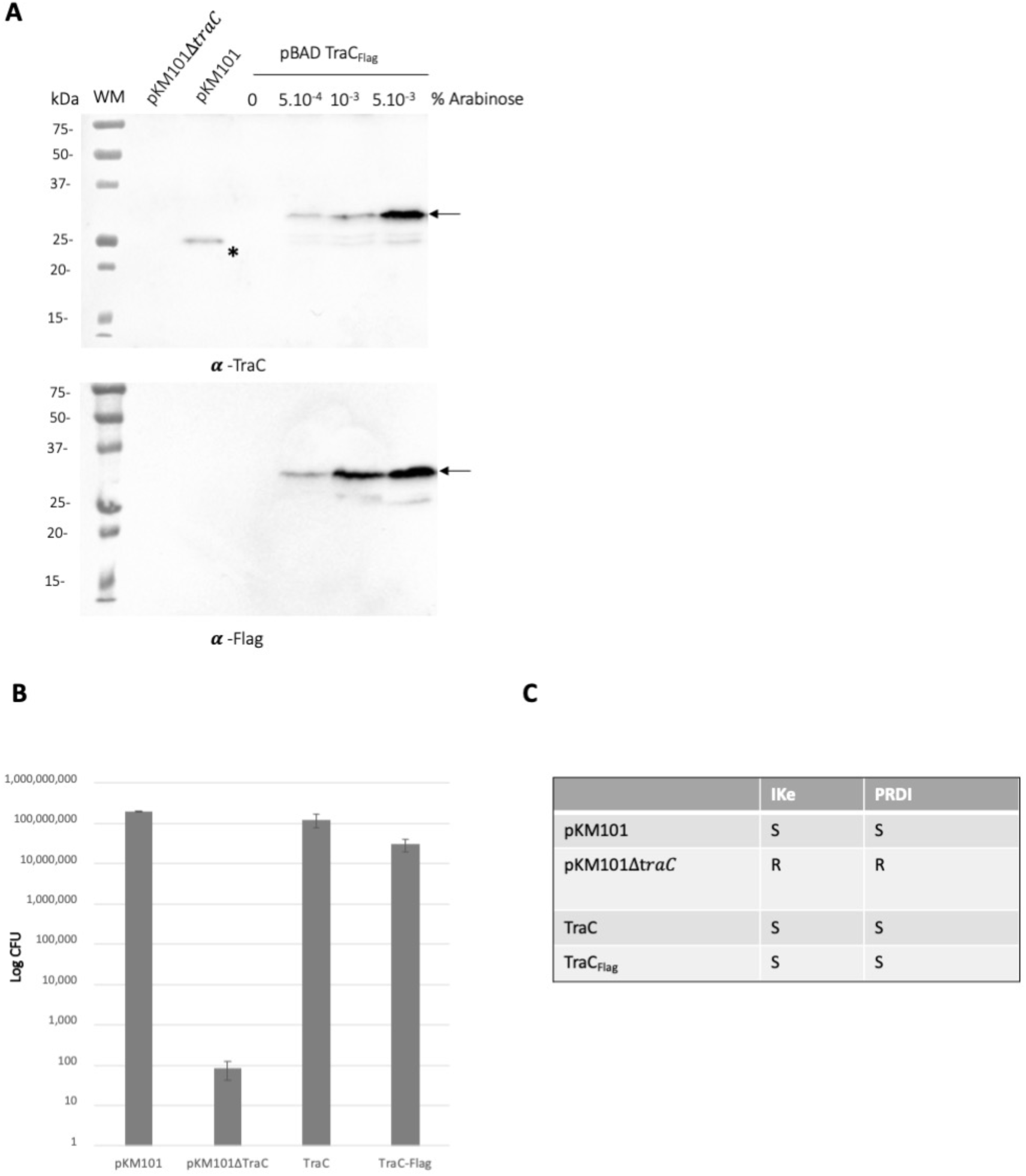
Functionality assays of TraC_Flag_. *(A)* Western blot analysis using TraC-specific antiserum and anti-Flag antibodies of TraC expression level in total cell lysate under native promotor or *ara* pBAD promoter using increasing level of induction by arabinose. Arrows indicate the molecular weight of TraC_Flag_; asterisk (*) indicate TraC. Functionality of TraC_Flag_ was monitored by conjugation (B) and phage infection (C) using 10^−3^% of arabinose. Values and S.D. (*error bars*) were calculated from three independent experiments.

**Supplementary Figure 4.**
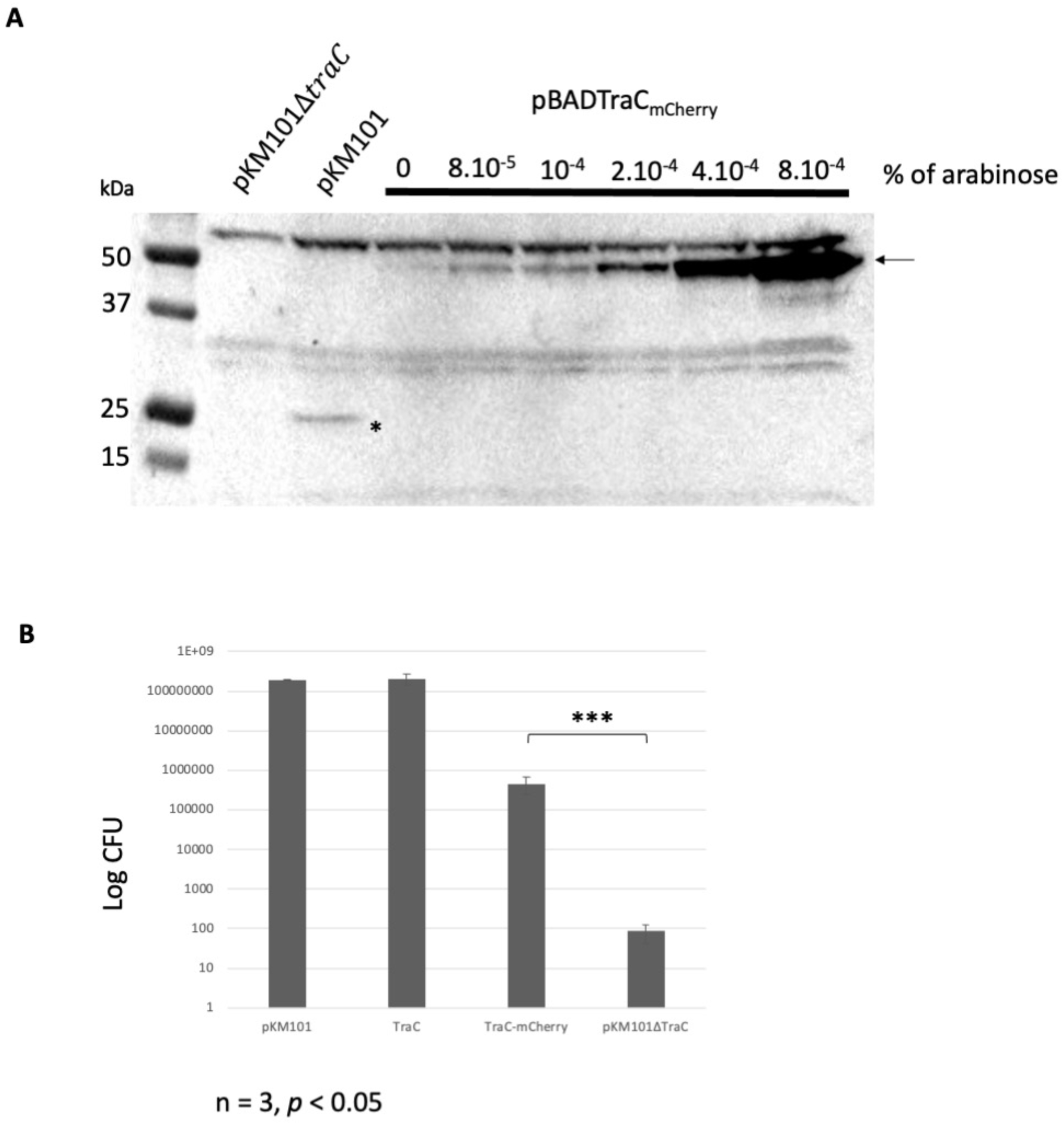
Functionality assays of TraC_mCherry_. *(A)* Western blot analysis using TraC-specific antiserum, of TraC_mCherry_ expression level in total cell lysate under *ara* pBAD promoter using increasing amount of arabinose for induction. *(B)* Functionality of TraC_mCherry_ was monitored by conjugation using 10^−4^% of arabinose. Arrow indicates TraC-mCherry; asterisk (*) indicate TraC.

